# A prevalent focused human antibody response to the influenza hemagglutinin head interface

**DOI:** 10.1101/2020.12.28.424595

**Authors:** Kevin R. McCarthy, Jiwon Lee, Akiko Watanabe, Masayuki Kuraoka, Lindsey R. Robinson-McCarthy, George Georgiou, Garnett Kelsoe, Stephen C. Harrison

**Affiliations:** Laboratory of Molecular Medicine, Boston Children’s Hospital, Boston, MA, 02115, USA; Harvard Medical School, Boston, MA, 02115, USA; Thayer School of Engineering, Dartmouth College, Hannover, NH, USA; Department of Immunology, Duke University, Durham, NC 27710, USA; Departments of Chemical Engineering, Molecular Biosciences, Biomedical Engineering, and Oncology, University of Texas at Austin, Austin, TX 78712, USA; Howard Hughes Medical Institute, Boston, MA 02115, USA

## Abstract

Novel animal influenza viruses emerge, initiate pandemics and become endemic seasonal variants that have evolved to escape from prevalent herd immunity. These processes often outpace vaccine-elicited protection. Focusing immune responses on conserved epitopes may impart durable immunity. We describe a focused, protective antibody response, abundant in memory and serum repertoires, to a conserved region at the influenza hemagglutinin head interface. Structures of eleven examples, eight reported here, from seven human donors demonstrate the convergence of responses on a single epitope. The eleven are genetically diverse, with one class having a common, IGκV1-39, light chain. All of the antibodies bind HAs from multiple serotypes. The lack of apparent genetic restriction and potential for elicitation by more than one serotype may explain their abundance. We define the head interface as a major target of broadly protective antibodies with the potential to influence the outcomes of influenza infection.

## INTRODUCTION

Influenza viruses threaten human health as both endemic (seasonal flu) and emerging (pandemic flu) pathogens. Four distinct seasonal influenza virus subtypes currently circulate each flu season. Each subtype evolves to escape dominant protective herd immunity elicited by past influenza exposures -- a process known as antigenic drift. Antigenically diverse influenza viruses emerge from animal reservoirs to initiate pandemics that sweep through immune naive human populations; between 1918 and 2009 this “antigenic shift” has occurred at least four times. Improved influenza vaccines will need to impart both seasonal and pre-pandemic immunity.

Focusing antibody responses on conserved epitopes is at the foundation of efforts to make improved or universal flu vaccine immunogens. Known broadly protective antibodies engage sites on the major virion surface protein, influenza hemagglutinin (HA). These include the receptor binding site (RBS) on the “head” of an HA subunit^1–5^, sites on the HA “stem”^6,7^, and a recently identified site at the interface between two heads of an HA trimer^4,8–10^. Antibodies to the RBS are potently neutralizing. Importance of residues at the periphery of the RBS for defining antigenic clusters suggests, however, that the most common of the RBS-directed antibodies are sensitive to the identity of those residues, probably limiting the long-term breadth of any such antibodies with footprints that extend beyond the sialic-acid pocket^11^. Stem and head-interface antibodies realize their full protective potential through Fc-dependent pathways that lead to killing of infected cells, rather than by blocking viral entry^12,13^. Broadly protective antibodies directed at these epitopes are likely present in most flu-exposed humans^3,5,9,14–16^.

We find that seasonal flu vaccination elicits a particularly strong serum antibody and memory B cell responses directed at the HA head interface. We describe here the anatomy of this dominant and focused response, by examining HA-Fab structures of eleven antibodies (including eight reported here) isolated from seven human subjects. The head interface accommodates contacts from structurally, and correspondingly genetically diverse, antibodies. A distinct subset (five antibodies from five different subjects) has a light-chain variable region encoded by the IgκV1-39 gene. Members of this germline-restricted subset bind HA nearly identically, and we define from structures the molecular basis for this restriction. All of the characterized antibodies bind to multiple HA serotypes, suggesting that the epitope may elicit an intrinsically broad response. These qualities of the head interface epitope can inform the design of improved influenza vaccines.

## RESULTS

### Origins of antibodies in this study

Structures of three HA-bound head-interface antibodies (H2214, S5V2-29 and FluA-20), from three different donors (designated EI-13, S5 and FluA-20 donor), have been reported previously^9,10,17^. We have now characterized six additional head-interface antibodies from two further donors (S1 and S8), identified by antibody/Fab competition. Antibodies D1 H1-3/H3-3, D1 H1-17/H3-14 and D2 H1-1/H3-1 were isolated from donors designated D1 and D2, respectively, and described originally as engaging the opposite surface of the contact between heads^4^. We determined that these antibodies instead compete for binding with a H2214 Fab on monomeric HA heads (Figure S1A). Lineage analysis^18^ suggests that both antibodies from D1 have arisen from a single naive progenitor. We have therefore assembled a panel of 11 unrelated antibodies, from seven donors, representing nearly all described examples of head-interface antibodies.

These antibodies arose from several different combinations of V, D, and J gene segments (Figure S1B). Combinations of nine V_H_, all from the IGHV-3 and IGHV-4 gene families, six D_H_ and four J_H_ gene segments encoded the eleven heavy-chain variable domains. HCDR3 lengths ranged from 9 to 25 amino-acid residues. Six different κ_V_ and four κ_J_ gene segments (among which, 2*01 appeared five times) encoded the eleven light chains, all V-kappa. In five of the eleven, IGκV1-39 had recombined with four different κ_J_ gene segments. LCDR3 lengths were between 5 and 10 residues.

### Structures

Structures of HA bound Fab fragments of S5V2-29, H2214 and FluA-20 have been reported^9^. We determined eight additional structures of Fab fragments from S1V2-51, S1V2-58, S1V2-83, S8V2-18, S8V2-37, S8V2-47, D1 H1-17/H3-14 and D2 H1-1/H3-1 in complex with various HA head domains (Figures 1A and B). All engage the same lateral surface of the HA head, although they do so at various angles and in various orientations with reference to the heavy and light chains. Despite the diversity of paratopes, all 11 Fabs ultimately converge upon a common site (Figure 1C). The five antibodies with light chains encoded by the IGκV1-39 gene segment, each from a different donor, all engage HA similarly (Figure 1B).

**Figure 1:**
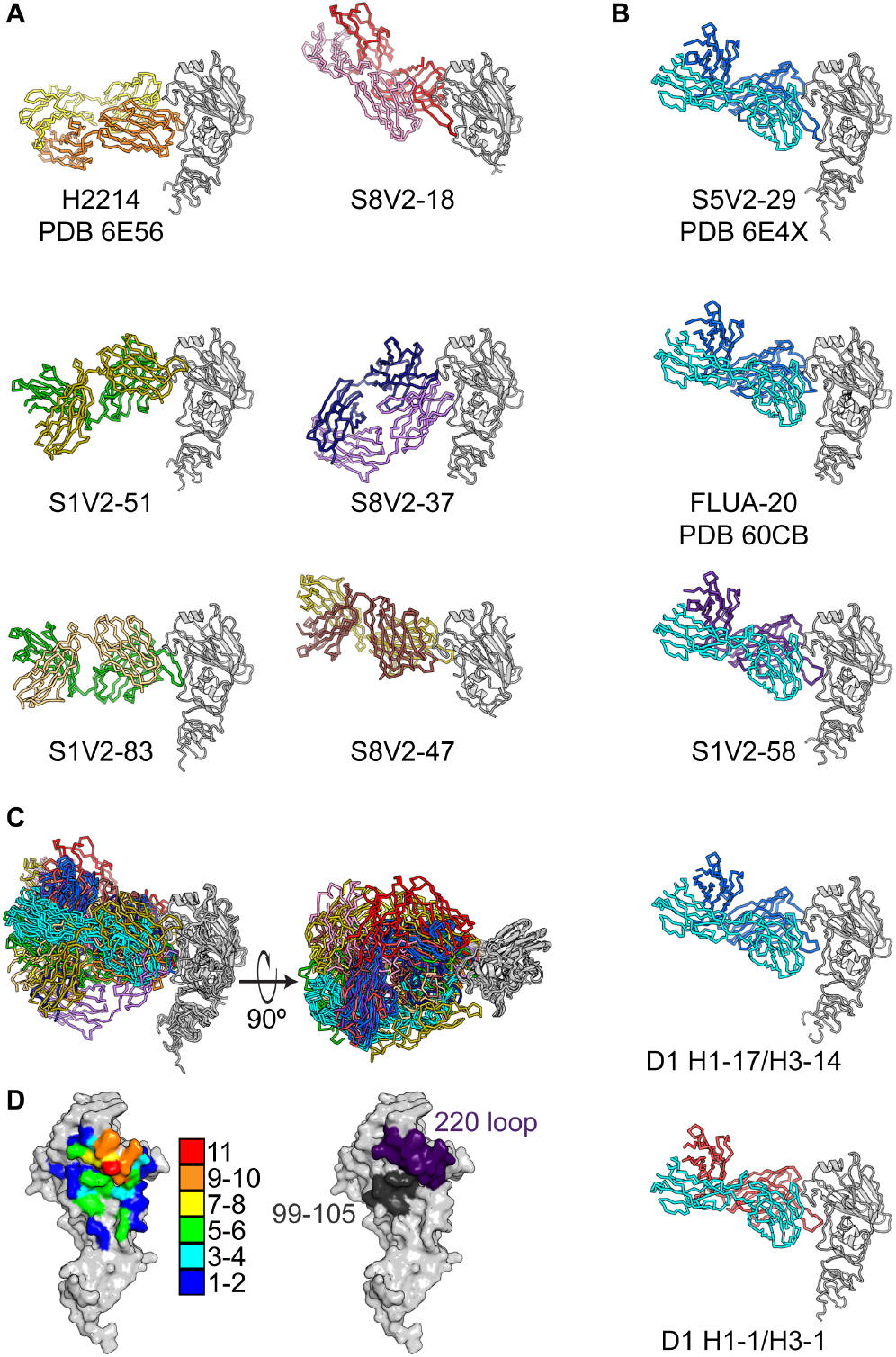
Structures of head interface antibodies bound to HA head domains. Fabs are colored according to the V_H_ (bolder color) and V_L_ (lighter color) gene usage. HA heads are all shown in gray. H2214, S5V2-29 and FluA-20 have been previously reported and their PDB IDs are indicated^9,10^. **A**. The six antibodies (two columns of three) that do not use the IGκV1-39 gene. **B**. One column showing the five antibodies that use IGκV1-39 (cyan). **C**. A superposition on the HA head domain of all 11 structures shown from two views. **D**. (Left) A contact heat map on the HA head surface. The number of Fabs (from a total of 11) contacting each residue were tallied and colored according to the key. (Right) Two key points of contact, the 220-loop (purple) and residues 99-105 (dark gray) are shown on the HA surface.

We have used these 11 structures to define the common head-interface epitope. The heat map in Fig. 1D illustrates a focused response, centered on the HA-220 loop (Figure 1D). All 11 antibodies contact Pro-221, and 10 of the 11 contact residues 222, 223 and 229. Other positions within this loop have contacts from more than half of the antibodies. The second most frequently contacted segment includes HA residues 99-105. Less frequently contacted sites ring these two “hot spots”.

### IGκV1-39 germline bias

Among the five IGκV1-39 encoded antibodies, four have heavy chains encoded by genes recombined from IGHV4-61 or IGHV4-59 and one, D2 H1-1/H3-1, from IGHV3-7. These antibodies present HA with a similar paratope. The HCDR3s, while unique in sequence and length, all curl towards the light chain and create a blunt antigen combining site that brings light chain framework regions 2 and 3 into contact with HA (Figure 2 and S1B). This arrangement accommodates the HA-220 loop in a groove at the heavy-light chain interface, with HCDR3 tucked beneath it.

**Figure 2:**
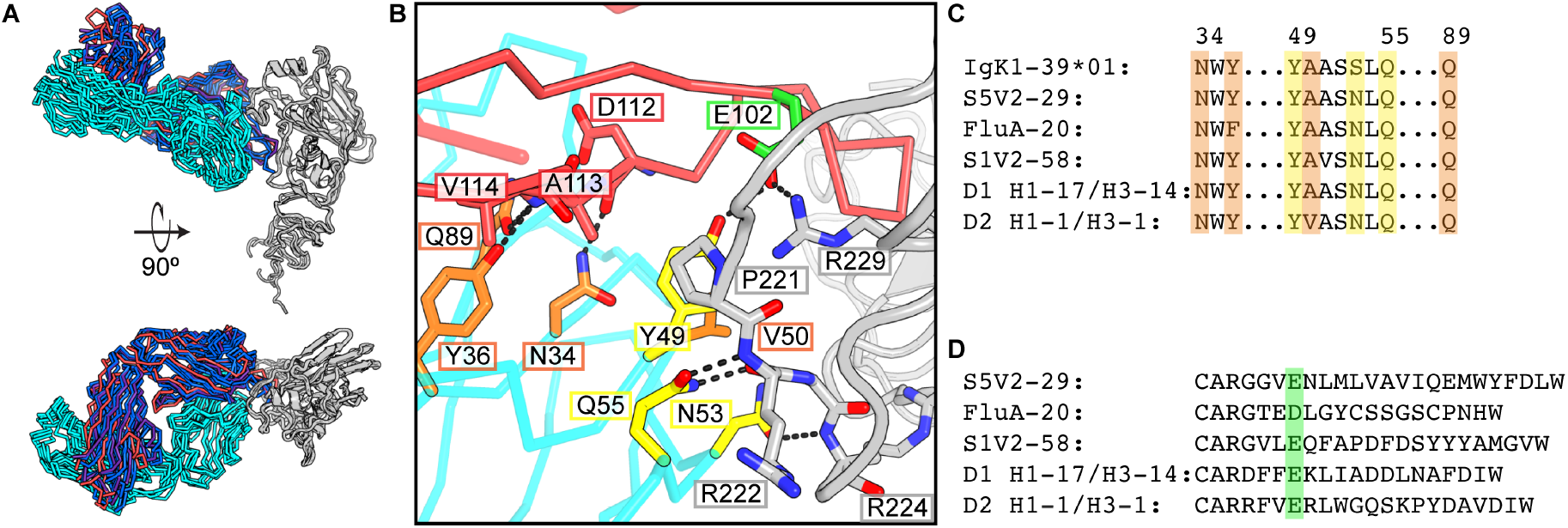
Stereotyped HA engagement by IGκV1-39 antibodies. **A.** Superposition on the HA head domain of the five IGκV1-39 antibodies. The HA heads are in gray, IGκV1-39, in cyan, and heavy chains are colored according to V_H_ gene usage and are the same as in Figure. 1. **B.** A zoomed in view of the IGκV1-39 interaction of D2 H1-1 H3-1 with HA from the bottom view of **A**. Key residues are shown in sticks. Light chain residues that contact HA are colored yellow and those that buttress/permit a curled HCDR3 conformation are colored in orange. A common acidic residue in HCDR3 is highlighted in green and shown in sticks. **C**. Partial sequence alignment of the germline IGκV1-39 sequence and the five antibodies. Shading is colored as in **B. D.** Sequence alignment of the antibody HCDR3s. An acidic residue at the sixth position of HCDR3 is shaded in green.

From these structures and the sequences of the human light chain V-gene repertoire, we can infer an IGκV1-39 signature to explain the stereotyped HA interaction. The principal contacts between the light chain and HA are from residues 49-55 -- YAASSLQ in the germline sequence (Figures 2B and S2). In all five structures, the side-chain of Gln at position 55 hydrogen bonds with both the main chain NH and the main-chain CO of HA residue 222. Also in all five, a Ser53Asn substitution allows the Asn side-chain to accept a hydrogen bond from the main-chain NH of HA residue 224, while donating a branched hydrogen bond to the carbonyls of residues 49 and 50 in the ß-turn at the tip of the LCDR2 loop. The side chain of Tyr49, anchored by a hydrogen bond with an acidic residue at the sixth position in HCDR3 in all five antibodies (Figure 2), is in van der Waals contact with conserved HA-Pro221. The HCDR3 acidic residue (supplied by IGHD3-3*01 or by n-nucleotides) then forms a double salt bridge with conserved HA-Arg229. The Tyr49-HCDR3-Asp/Glu interaction and Tyr49-HCDR3 van der Waals interactions buttress the curl of HCDR3. The small amino-acid residue at light chain position 50 avoids clashes with HCDR3 and HA, while making van der Waals contacts with each. Additional HCDR3 stabilization appears to come from main-chain hydrogen bonds with side chains of light chain residues Asn34, Tyr36 (Phe in FluA20, perhaps dispensable after affinity maturation), and Gln89. The motif of 49-Y-small-X-X-N-X-Q-55, Asn34, Tyr36, Gln89 is unique to IGκV1-39 (Figure 2 B-D and S2). Because much of this motif is germline encoded it also likely explains the abundance of this gene segment among those encoding the light chains of these antibodies.

### Unique examples

The six antibodies with other light chains are quite diverse in the ways in which they contact HA (Figure 3 A and B). For those antibodies, gene usage does not correlate with HA contacts. As an ensemble they represent the products of multiple alternative affinity maturation pathways for head interface engagement. We describe below the five newly reported here.

**Figure 3:**
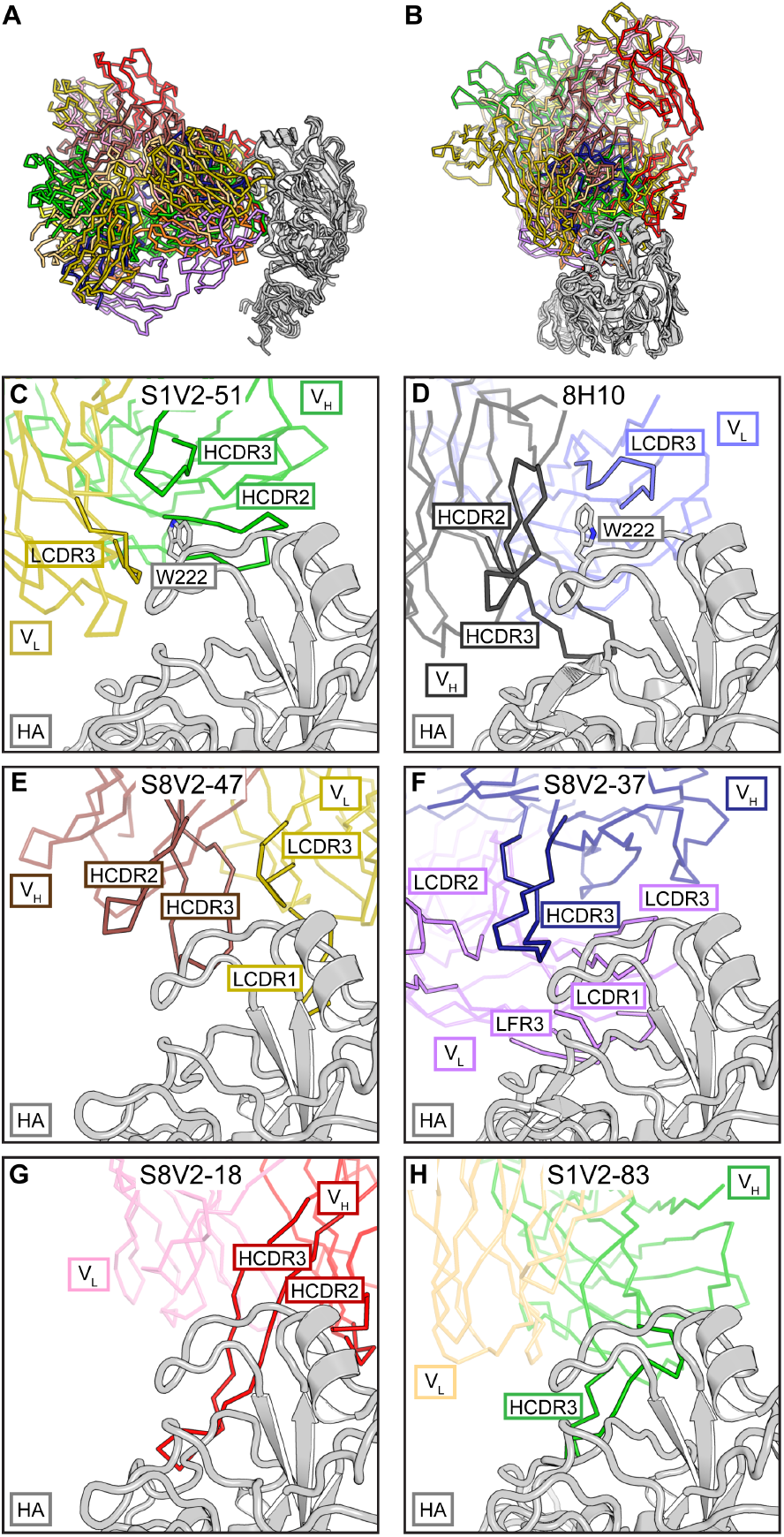
Diversity of non-IGκV1-39 head interface antibodies. **A.** All six structures are shown with a common orientation for the HA head domain. Fabs are colored according to V_H_ (bolder color) and V_L_ (lighter color) gene usage; HA heads are in gray. **B.** Top views of the structures in **A**. Panels **Cs-H** are details of this view. Fab fragments are faded except for selected CDR or framework regions (FR) to highlight the diversity of Fab-HA contacts. **C.** S1V2-51. HA residue Trp-222 is coordinated in a hydrophobic vault by HCDRs 2 and 3 and LCDR3 that is similar to mouse antibody 8H10. **D.** 8H10^8^ (PDB 6N5B) and its interaction with Trp-222. **E.** S8V2-47. HCDRs 2 and 3 and LCDRs 1 and 3 all contribute to the interaction with HA. **F.** S8V2-37. HCDR3 displaces LCDR2 to create an intertwined antigen combining site with HCDR3 and LCDRS1,2 and LFR3. **G.** S8V2-18. Among the interface antibodies HCDRs 2 and 3 contact the most recessed surfaces of the HA head. **H.** S1V2-83. Most of the interaction with HA is through HCDR3.

S1V2-51 has a short, nine amino acid HCDR3 that lies within the heavy-light chain interface and creates a hydrophobic vault, into which HA Pro221 and Trp222 insert (Figure 3C). HCDR3 caps the vault and supplies polar residues including Asp99 that forms a salt bridge to the Nε1 of T rp222. HA engagement is similar to murine interface antibodies that also coordinate Pro221 and Trp222 within the hydrophobic heavy-light chain interface (Figure 2D). Human and mouse antibodies also contact residues of the receptor binding site. In S1V2-51 a Phe extends from a long CDRL1 to make van der Walls contacts with conserved sialic acid coordinating residue Leu226. S8V2-47 uses the same light chain V segment as S1V2-51, IGkV4-1*01. In these examples HA binding is not biased by gene usage as their light chains engage opposite sides of the head interface epitope (Figure 3C and 3E). In S8V2-47 the HA 220-loop, including Pro221 is pinched between HCDRs 2 and 3 (Figure 3E).

The approach of S8V2-37 is substantially different from those of the other interface antibodies, as it engages HA with a distinctive antigen combining site. HCDR3 has displaced LCDR2 and flanking parts of framework regions 2 and 3, creating an intertwined paratope (Figure 3F). Contacts with HA are nearly all through HCDR3, LCDR1, LCDR3 and light chain framework 3.

S8V2-18 contacts HA almost entirely through its heavy chain, due to a long HCDR3. The light chain contributes one van der Walls interaction with HA-Pro221. S8V2-18 contacts the most recessed surfaces of any of the interface antibodies. Its heavy chain hooks around the HA head and contacts residues facing the 3fold axis of the trimer (Figure 3G). S1V2-83, has the longest HCDR3 (l25-amino acids) which adopts a Γ-shaped conformation. The arm contacts conserved sites on the HA-220 loop and its stem extending downward along the interface epitope (Figure 3H). This interaction contributes most of the contacts with HA. Both S1V2-83 and S1V2-51 are both V(D)J recombinants derived from IGH3-30*01; this gene usage does not appear to impose a biased mode of binding (Figure 3C and 3H).

### Broad HA reactivity of interface antibodies

We examined by ELISA the binding of these antibodies to divergent HA subtypes (Figure 4 A and B). All of them bound HAs from at least two serotypes, and all but one bound at least one group 1 and group 2 HA. The antibodies belonging to the IGκV1-39 subclass had the greatest breadth of binding. Three bound to 10 and one bound to 11 of the 12 HA serotypes assayed. The combined breadth of the subject 8 antibodies, without an IGκV1-39 representative, covered 10 of the 12 HA serotypes, demonstrating that polyclonality and diverse V(D)J gene usage can achieve broad reactivity.

**Figure 4:**
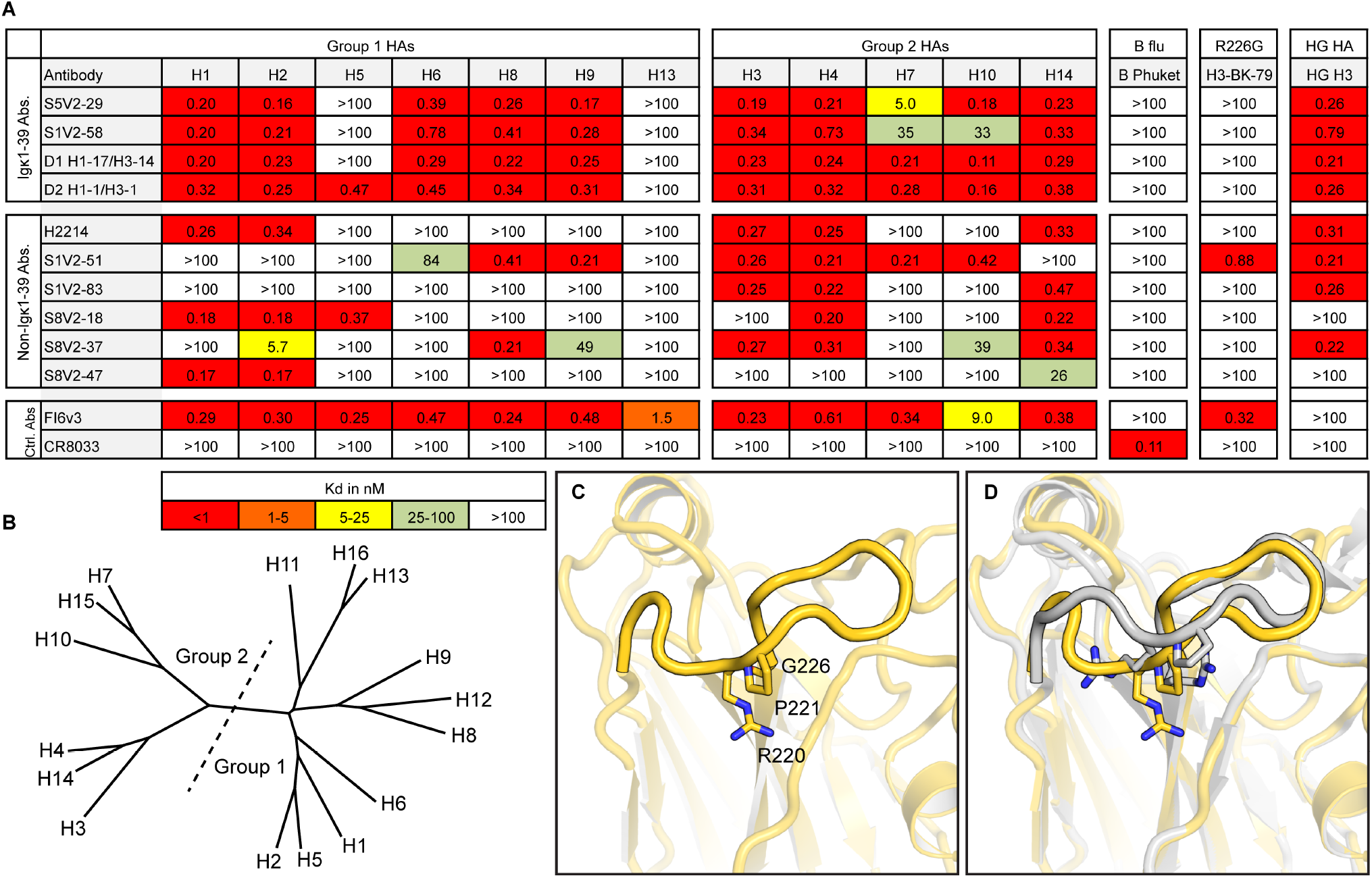
Breadth of binding by head interface antibodies. **A.** Dissociation constants from ELISA measurements. The pan-influenza A binding, stem-directed antibody, FI6v3^12^, was used as a positive control and an influenza B-specific, RBS-directed antibody, CR8033^31^, as a negative control for influenza A isolates. Boxes are colored according to the key at the bottom. HAs used: H1 A/California/04/2009(H1N1, H2 A/Japan/305/1957(H2N2), H3 A/Hong Kong/JY2/1968(H3N2), H4 A/American black duck/New Brunswick/00464/2010(H4N6), H5 A/Viet Nam/1203/2004(H5N1), H6 A/Taiwan/02/2013(H6N1), H7 A/Taiwan/01/2017(H7N9), H8 A/northern shoveler/California/HKWF1204/2007(H8N4), H9 A/Beijing/1/2017(H9N2), H10 A/Jiangxi/IPB13/2013(H10N8), H13 A/gull/Maryland/704/1977(H13N6), A/mallard/Wisconsin/10OS3941/2010(H14N6), B-Phuket B/Phuket/3073/2013, H3-BK-79 A/Bangkok/01/1979(H3N2), gHA^shield 8^. **B**. Phylogram of 16 HA serotypes. The division of HA groups 1 and 2 are denoted with a dotted line. **C.** Structure of the 220-loop of A/Bangkok/01/1979(H3N2). The HA head is faded except for the 220-loop to emphasize this feature. Residues Arg220, Pro221, Gly229 are shown in sticks. **D.** Comparison with A/Aichi/02/1968(H3N2) (PDB ID: 2VIU) (gray) which has an Arg at position 229.

The polyclonality of the head interface response should, in principle, prevent pan-virus resistance. Among circulating strains of most HA serotypes, Arg229 is maintained by strong purifying selection. Some laboratory passaged viruses have a substitution for this residue^9^. Most head interface antibodies contact Arg229 (Figure 1D) and fail to bind 229 mutants (Figure 4A). We determined the HA trimer structure of laboratory propagated A/Bangkok/01/1979(H3N2) with its acquired Arg229Gly substitution. The mutation led to distortion of the 220-loop, the major contact site for all head interface antibodies (Figures 1D, 4C, 4D). The reconfigured loop in our structure was unlike that in other HAs in the PDB (aside from one other R229 mutant, PDB ID 6MXU). Despite the substitution and rearrangement, S1V2-51 bound the A/Bangkok/01/1979(H3N2) isolate (Figure 4A). Constraints on the 220-loop, which forms one side of the receptor-binding pocket, are likely to limit potential pathways of for antibody escape.

We also assayed binding to a hyperglycosylated HA immunogen, gHA^shield 8^ based on an A/Aichi/02/1968(X-31)(H3N2) template, (Figure 4A) Mice immunized with gHA^shield^ have higher frequencies of head interface antibodies than do those immunized with WT^8^. All of the human antibodies that bind wild type A/Aichi/02/1968(X31)(H3N2) (n=8) also bind gHA^shield^ with comparable affinity. Similar immunogens may boost or selectively elicit desired classes of head interface antibodies in humans.

## DISCUSSION

Influenza HA head-interface directed, circulating memory B cells are abundant in humans following flu vaccination, as are interface-directed serum antibodies. In donors S1, S5 and S8, ~4% of the influenza A reactive memory B-cell repertoire encoded interface-directed, B-cell receptors^9^. Serum antibodies from donors D1 and D2, were quantified by mass spectrometry^4^. In the H1-directed response, antibody D2 H1-1/H3-1 ranked first and accounted for >15% of the influenza A antibody repertoire at day 28 post vaccination. The two D1 antibodies (H1-3/H3-3 and H1-17/H3-14) appear to be clonally related; H1-3/H3-3 is the third most abundant antibody in the vaccine responses to both the H1 and H3 components, together accounting for about 11% of the influenza A antibody repertoire at day 28. These antibodies were also present pre-vaccination, reflecting the abundance of cells with the same clonotype in the long-lived plasma-cell compartment^4^. Thus, the influenza HA head interface is immunogenic, and it elicits a strong serum response and a robust seeding of B cell memory. The structures described here define the anatomy of this dominant, focused antibody response to a single epitopic surface.

Contacts with HA from the eleven antibodies (eight newly reported here) from seven human donors overlap each other substantially and define the core epitope. They show that structurally and genetically diverse antibodies can engage this core, while varying slightly in their interactions with its periphery. One antibody subclass has a mode of HA binding dictated by the IGκV1-39 gene. The five examples from five human donors bind nearly identically. Apparent determinants for this bias are largely germline encoded; they are unique to IGκV1-39 *01 (or an identical duplicated copy IGκV1-39 *01). Within the IGκV1-39 subclass of interface antibodies, common substitutions reflect convergent affinity maturation pathways, as does the presence of a D-gene or n-nucleotide encoded residue in HCDR3. The acquired substitutions, all at the same nucleotide, occur in an AID mutation hotspot, perhaps contributing to the abundance of this subclass (Figure S2).

Why is the interface epitope immunogenic at all, since it is largely occluded in the “ground state” conformation of the trimer? The failure of interface-directed antibodies to neutralize in single-round infectivity assays suggests that, like the HA stem, the epitope is poorly accessible on virions^8–10^. Nonetheless, members of both these classes of non-neutralizing antibodies protect mice from lethal challenge by Fc-mediated mechanisms, implying that the corresponding epitopes are more readily accessible when HA is present on the cell surface. Since all but one of the samples from which the antibodies in this study were obtained were taken following immunization with TIV or QIV (and hence exposure to isolated protein, rather than intact virions), conformational fluctuations in the split-vaccine HA immunogen would probably have been at least as pronounced as it is for HA on the surface of a cell. Moreover, our incomplete understanding of how and in what form antigen is presented to B cells by follicular dendritic cells in germinal centers leaves open additional mechanisms for the strength of a secondary response to one epitope with respect to another.

Conservation of the head interface may also contribute to its apparent immunodominance. Imprinting by an initial exposure appears to condition all later response to influenza HA. A conserved epitope, even if only transiently exposed, may become immunodominant “by default” if most other immunogenic epitopes have mutated. For example, conservation in most H1 isolates since 1918 of a lateral patch on the head of H1 HAs caused this epitope to dominate humoral responses to the 2009 pandemic H1, which differed on other exposed surfaces from the variants most living individuals would have seen^19,20^. Unlike the head interface, the lateral patch has no apparent structural or functional role, and mutations quickly appeared in circulating HAs, leading to prevalence of the resistant viruses within 2-3 years^19^. Thus, even an initially subdominant epitope can become dominant if it is the only one well represented in the memory compartment.

Stem-directed antibodies appear to be less abundant than interface-directed ones, despite broad conservation of the relevant surface. As we have shown here, the latter have diverse germline origins and few paratopic constraints, while the former have constrained gene usage and limited paratopic diversity^14–16^. The number of naive, mature B cells in a population expressing randomly recombined BCRs should therefore be much higher for the genetically diverse interface response than for the more restricted stem response. Moreover, the frequency of H1-H3 cross reactive interface antibodies implies that imprinting by either an H1 or an H3 subtype would establish a common memory pool. For the head interface directed antibodies, V_H_ mutation frequencies ranged between 3.5-10%, consistent with recall responses and comparable to the levels observed for stem directed antibodies.

A major open question is whether antibodies to the stem or interface afford protection or reduce the severity of infection in humans. Preexisting, but incomplete, immunity to influenza can lessen disease severity and duration, as documented for flu seasons in which circulating and vaccine isolates are antigenically mismatched^21–23^; effector functions of non-neutralizing antibodies might account for some of this reduction. For conserved surfaces such as the head interface, such antibodies would also confer some degree of pre-pandemic protection^9,10^. Vaccines that elicit disease-burden limiting immunity might thus prove valuable for both recurrent seasonal infections and for emerging viruses.

## METHODS

### Cell lines

Human 293F cells were maintained at 37°C with 5% CO2 in FreeStyle 293 Expression Medium (ThermoFisher) supplemented with penicillin and streptomycin. High Five™ Cells (BTI-TN-5B1-4) (Trichoplusia ni) were maintained at 28°C in EX-CELL 405 medium (Sigma) supplemented with penicillin and streptomycin.

### Recombinant Fab expression and purification

Synthetic heavy- and light-chain variable domain genes for Fabs were cloned into a modified pVRC8400 expression vector, as previously described^5,24,25^. Fab fragments used in crystallization were produced with a C-terminal, noncleavable 6xhistidine (6xHis) tag. Fab fragments used for binding studies were cloned into a pVRC8400 vector that was further modified by the introduction of a rhinovirus 3C protease cleavage site between the heavy chain constant domain and a C-terminal 6xHis tag^5^ Fab fragments were produced by polyethylenimine (PEI) facilitated, transient transfection of 293F cells that were maintained in FreeStyle 293 Expression Medium. Transfection complexes were prepared in Opti-MEM and added to cells. Supernatants were harvested 4-5 days post transfection and clarified by low-speed centrifugation. Fabs were purified by passage over Co-NTA agarose (Clontech) followed by gel filtration chromatography on Superdex 200 (GE Healthcare) in 10 mM Tris-HCl, 150 mM NaCl at pH 7.5 (buffer A). For binding studies the 6xHis tag were removed from some Fabs by treatment with PreScission protease (MolBioTech; ThermoScientific) and the protein repurified on cobalt-nitrilotriacetic acid (Co-NTA) agarose (Clontech) followed by gel filtration chromatography on Superdex 200 (GE Healthcare) in buffer A to remove the protease, tag, and uncleaved protein.

### Recombinant IgG expression and purification

The heavy chain variable domains of selected antibodies were cloned into a modified pVRC8400 expression vector to produce a full length human IgG1 heavy chain. IgGs were produced by transient transfection of 293F cells as specified above. Five days post-transfection supernatants were harvested, clarified by low-speed centrifugation, and incubated overnight with Protein A Agarose Resin (GoldBio). The resin was collected in a chromatography column, washed with a column volume buffer A, and eluted in 0.1M Glycine (pH 2.5) which was immediately neutralized by 1M tris(hydroxymethyl)aminomethane (pH 8). Antibodies were then dialyzed against phosphate buffered saline (PBS) pH 7.4.

### Recombinant HA expression and purification

Recombinant (HA) constructs were expressed by infection of insect cells with recombinant baculovirus as previously described^1,9,24^.. In brief, a synthetic DNA corresponding to the full-length ectodomain (FLsE) or the globular HA-head was subcloned into a pFastBac vector modified to encode a C-terminal thrombin cleavage site, a T4 fibritin (foldon) trimerization tag, and a 6xHis tag. The resulting baculoviruses produce recombinant HA (rHA) trimers and trimeric HA heads. Monomeric HA-heads were produced by subcloning DNAs corresponding to the HA-head domain into a pFastBac vector modified to encode a C-terminal rhinovirus 3C protease site and a 6xHis tag. Supernatant from recombinant baculovirus infected High Five™ Cells (Trichoplusia ni) was harvested 72 hr post infection and clarified by centrifugation. Proteins were purified by adsorption to Co-NTA agarose resin, followed by a wash in buffer A, a second wash (trimers only) with buffer A plus 5-7mM imidazole, elution in buffer A plus 350mM imidazole (pH 8) and gel filtration chromatography on a Superdex 200 column (GE Healthcare) in buffer A.

gHA^shield 8^ was produced by polyethylenimine (PEI) facilitated, transient transfection of 293F cells maintained in FreeStyle 293 Expression Medium. Transfection complexes were prepared in Opti-MEM and added to cells. Supernatants were harvested 4-5 days post transfection and clarified by low-speed centrifugation. Fabs were purified by passage over Co-NTA agarose (Clontech) followed by gel filtration chromatography on Superdex 200 (GE Healthcare) in 10 mM Tris-HCl, 150 mM NaCl at pH 7.5 (buffer A).

### Biolayer interferometry (BLI)

Binding of Fabs with H_A_ heads was analyzed by BLI (BLItz: forteBIO; Pall); all measurements were in buffer A at room temperature. For competition assays purified His tagged Fabs were immobilized on a Ni-NTA biosensor. HA head domains and Fab fragments with the His tag removed were used as analytes.

### ELISA

Five hundred nanograms of rHA FLsE were adhered to high-capacity binding, 96 well-plates (Corning) overnight in PBS pH 7.4 at 4°C. Pates were blocked with PBS containing 2% bovine serum albumin (BSA) and 0.05% Tween-20 (PBS-T) for 1 hr at room temperature. Blocking solution was removed, plates were washed once with PBS-T, and 5-fold dilutions of IgGs (in PBS-T) were added to wells. Plates were then incubated for 1 hr at room temperature followed by removal of IgG solution and three washes with PBS-T. Secondary, antihuman IgG-HRP (Abcam ab9722) diluted 1:20,000 in PBS, in PBS-T was added to wells and incubated for 30 min at 37°C. Plates were then washed three times with PBS-T. Plates were developed using 1-Step ABTS substrate (ThermoFisher, Prod#37615). Following a brief incubation at room temperature, HRP reactions were stopped by the addition an equal volume of 1% sodium dodecyl sulfate (SDS) solution. Plates were read on a BioTek™ ELx808™ Microplate Reader at 405 nm.

### Crystallization

Fab fragments were co-concentrated with HA-head domains) at a molar ratio of ~1:1.3 to a final concentration of ~20 mg/ml. Crystals of Fab-head complexes were grown in hanging drops over reservoir solutions indicated in (Figure S3) Crystals were cryprotected in the well solution for crystals obtained in PEG 400 at concentrations ≥30%. Other crystals were cryprotected with glycerol at concentrations of 12-25% in cryoprotectant buffers that were 20% more concentrated than the well solution. Cryprotectant was added directly to the drop, crystals were harvested, and flash cooled in liquid nitrogen.

### Structure determination and refinement

We recorded diffraction data at the Advanced Photon Source on beamlines 24-ID-E and 24-ID-C. Data were processed with XDS^26^. Molecular replacement was carried out with PHASER^27^, dividing each complex into four search models (Figure S3). We carried out refinement calculations with PHENIX^28^ and model modifications, with COOT^29^. Refinement of atomic positions and B factors was followed by translationliberation-screw (TLS) parameterization and, if applicable, placement of water molecules. Final coordinates were validated with the MolProbity server^30^. Data collection and refinement statistics are in Table S4. Figures were made with PyMOL (Schrödinger, New York, NY).

## DATA AND SOFTWARE AVAILABILITY

Coordinates and diffraction data have been deposited at the PDB, accession numbers 6XP0, 6XPQ, 6XPR, 6XPX, 6XPY, 6XPZ, 6XQ0, 6XQ2 and 6XQ4.

## ACKNOWLEDGMENTS

We thank the many members of our Program Project Consortium for advice and discussion. X-ray diffraction data were recorded at beamline ID-24-E (operated by the Northeast Collaborative Access team: NE-CAT) at the Advanced Photon Source (APS, Argonne National Laboratory). We thank NE-CAT staff members for advice and assistance in data collection. NE-CAT is funded by NIH grant P30 GM124165. APS is operated for the DOE Office of Science by Argonne National Laboratory under contract DE-AC02-06CH11357. The research was supported by NIAID Program Project Grant P01 AI089618 (to S.C.H.).

## SUPPORTING FIGURES

**Figure S1:**
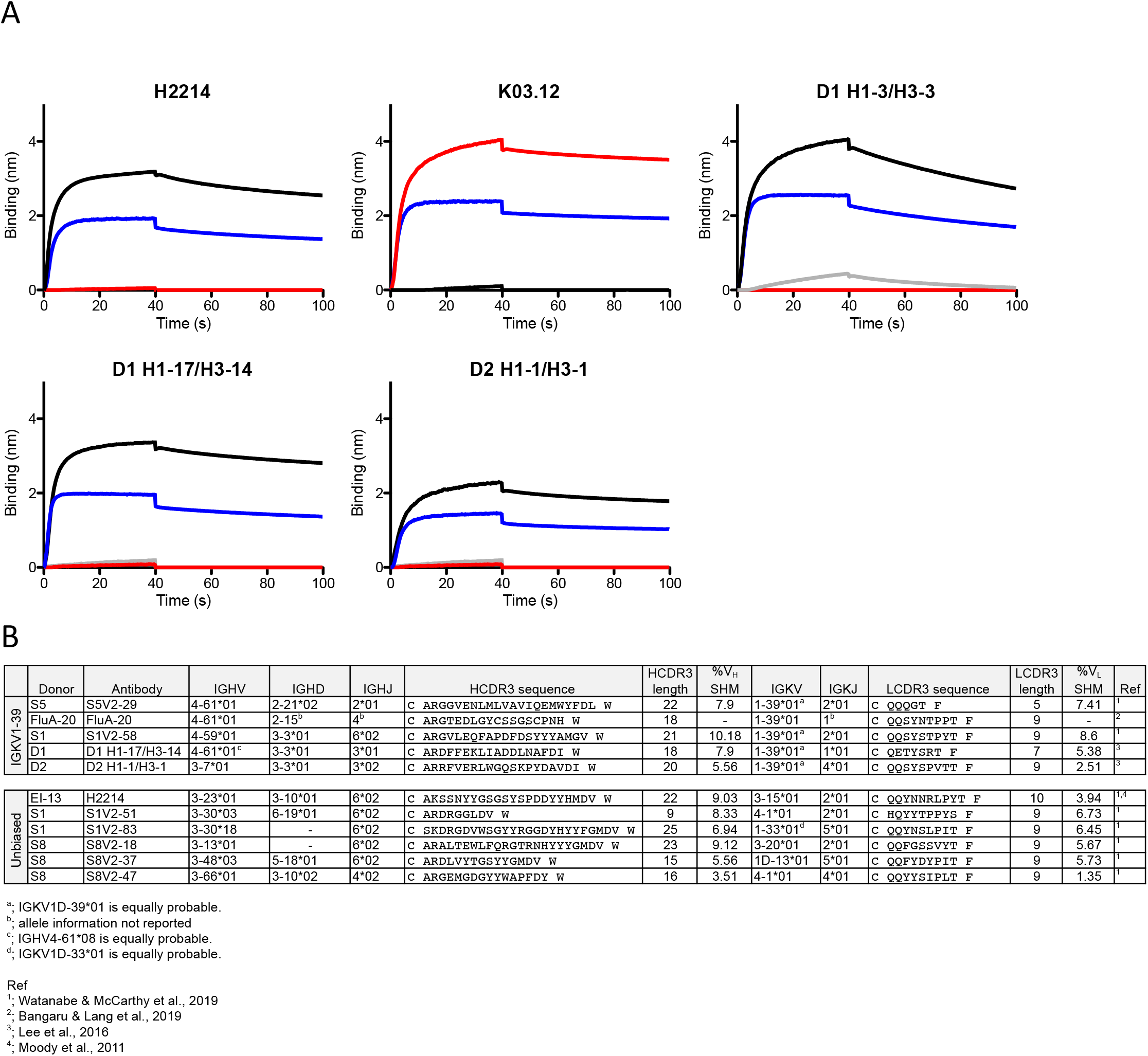
Head interface antibodies. **A.** Identification of additional head interface antibodies. Panels show traces for association and dissociation of an HA head domain of A/Texas/50/2012(H3N2), at a concentration of 12 μM, with the Fab fragment of the antibody shown in the panel header, immobilized on a BLI sensor. In each panel, the blue curve shows binding in the absence of any competitor; the red curve, in the presence of a 4-fold molar excess of H2214^9^ (head interface-directed), pre-incubated with the HA head; the black curve, in the presence of a 4-fold molar excess of Fab from an RBS-directed antibody K03.12^5^ pre-incubated with the HA head; the curve for D1 and D2 antibodies, competition with the antibody named in the panel header in the presence of a 4-fold molar excess of Fab (competition with itself). **B.** Gene usage, H/LCDR3 sequences and percent somatic hypermutation (SHM) of human HA head interface antibodies discussed in this manuscript.

**Figure S2:**
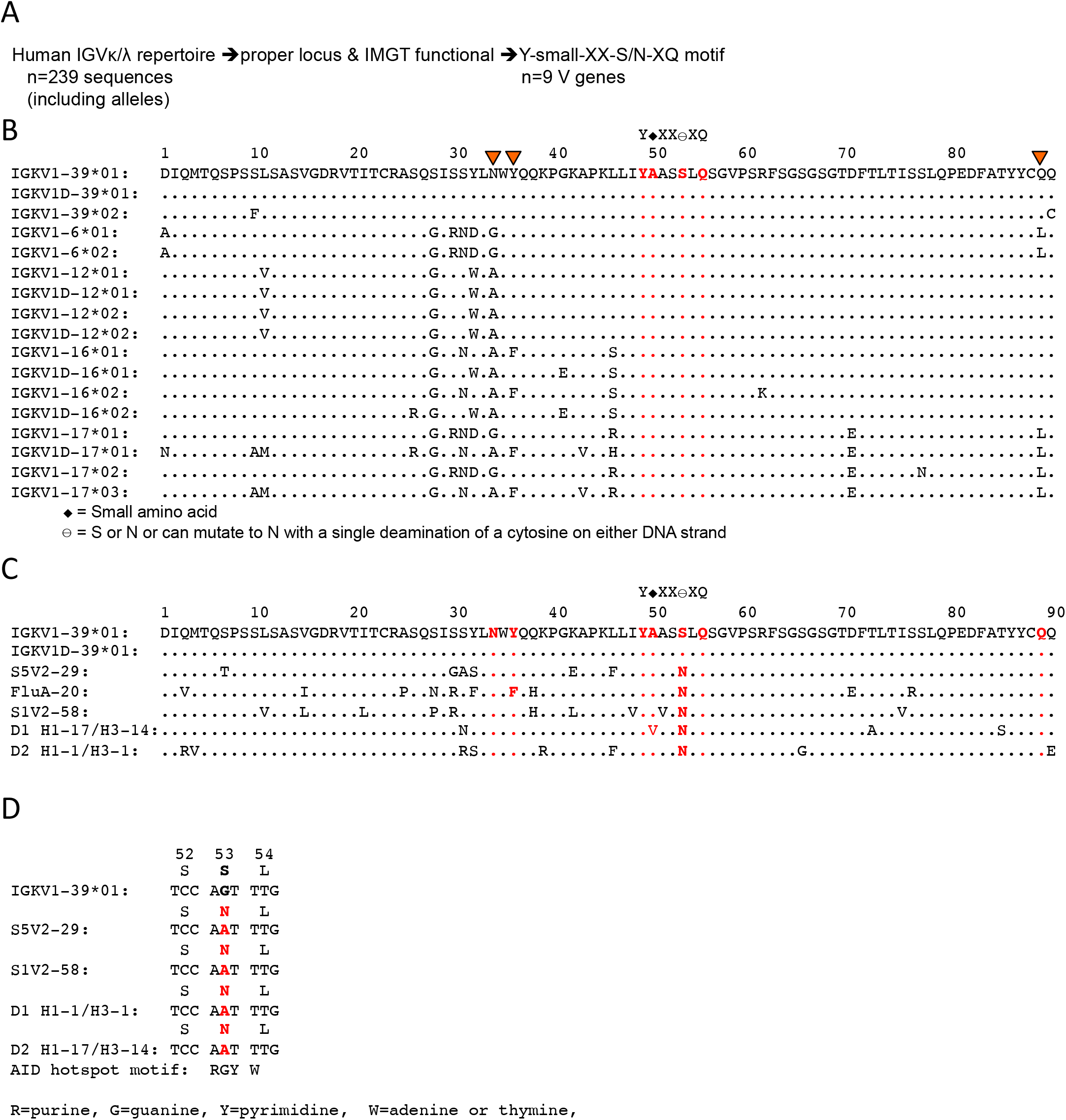
Identification of the genetic contribution for the IG κ V1-39 gene bias. **A.** Workflow using a motif search informed by structural analysis. **B.** Alignment of V genes that fulfill the sequence motif search. Residues in the motif care colored red. Residues that buttress HCDR3 are denoted with orange triangles. Only alleles or an identical duplication of IGκV1-39 have both the motif and buttressing residues. **C.** Sequence alignment of the germline IGκV1-39 and the IGκV1-39 antibodies that were characterized. Key residues are shown in red. **D.** Codon analysis of the S53N mutation that is present in all IGκV1-39 antibodies. Codons 52, 53 and 54 are shown. The S53N mutation is due to a single nucleotide change (colored red), introduced by AID. The cytosine occurs in an AID hotspot sequence motif.

**Figure S3:**
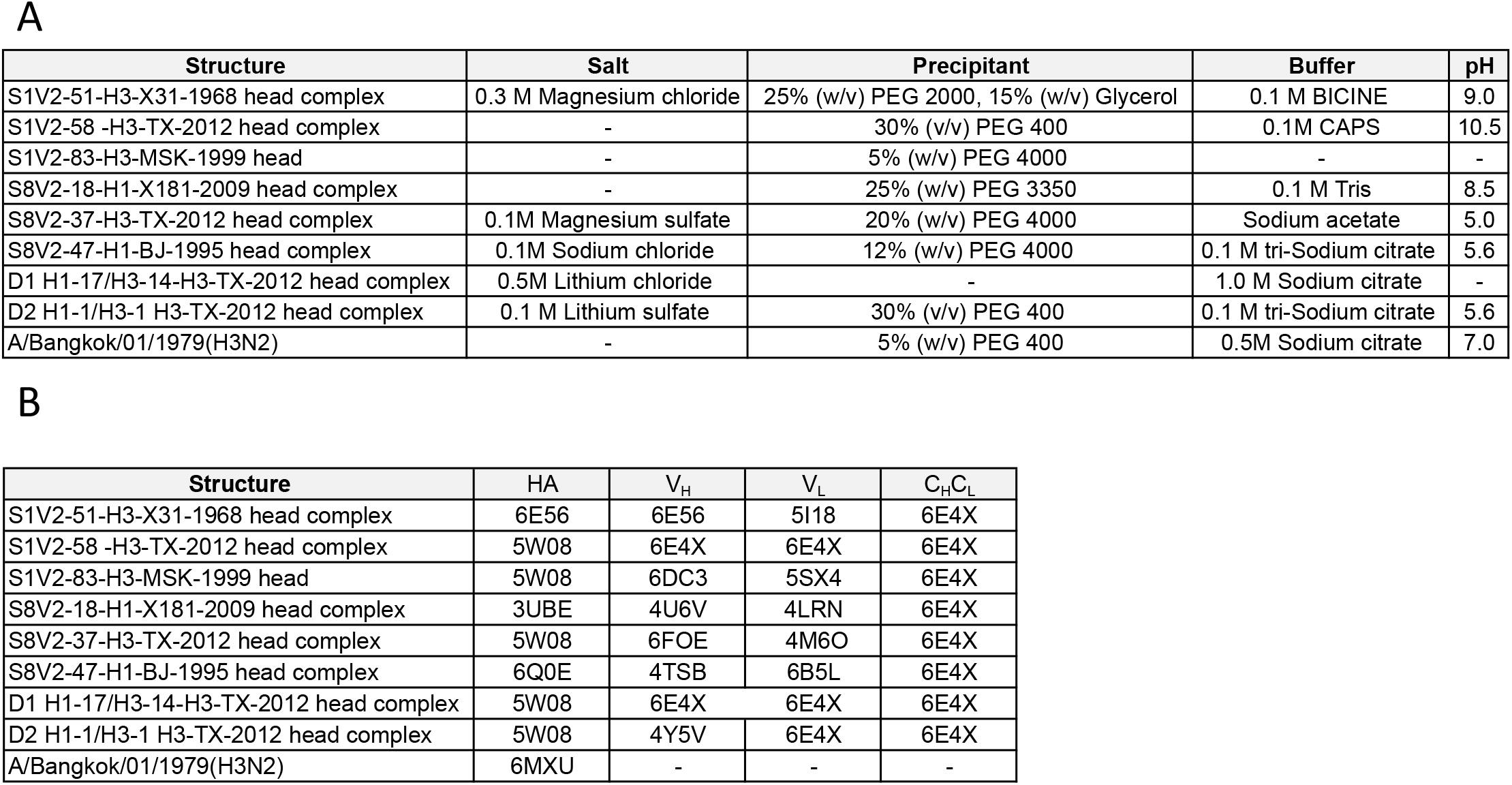
Summary of crystallization conditions and molecular replacement models.

**Table S1:**
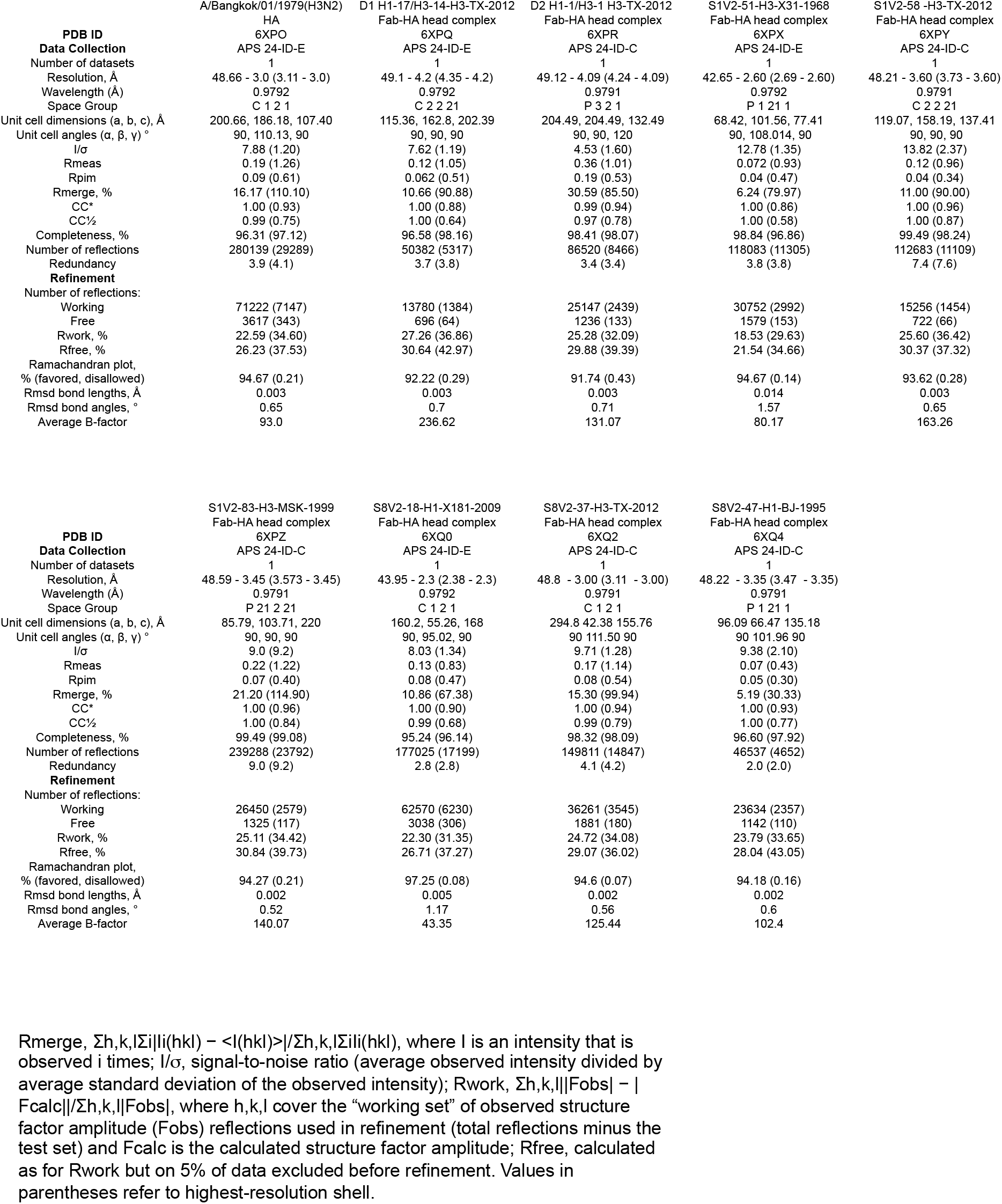
Data collection and refinement statistics.

## Notes

### Competing Interest Statement

The authors have declared no competing interest.

